# Quantifying antibiotic impact on within-host dynamics of extended-spectrum beta-lactamase resistance in hospitalized patients

**DOI:** 10.1101/548453

**Authors:** Rene Niehus, Esther van Kleef, Mo Yin, Agata Turlej-Rogacka, Christine Lammens, Yehuda Carmeli, Herman Goossens, Evelina Tacconelli, Biljana Carevic, Surbhi Malhotra-Kumar, Ben S Cooper

## Abstract

Antibiotic exposure can perturb the human gut microbiome and cause changes in the within-host abundance of the genetic determinants of drug-resistance in bacteria. Such within-host dynamics are expected to play an important role in mediating the relationship between antibiotic use and persistence of drug-resistance within a host and its prevalence within a population. Developing a quantitative representation of these within-host dynamics is an important step towards a detailed mechanistic understanding of the population-level processes by which antibiotics select for resistance. Here we study extended-spectrum beta-lactamase (ESBL) producing organisms of the Enterobacteriaceae bacterial family. These have been identified as a global public health priority and are resistant to most first-line antibiotics for treatment of Enterobacteriaceae infections.

We analyse data from 833 rectal swabs from a prospective longitudinal study in three European countries including 133 ESBL-positive hospitalised patients. Quantitative polymerase chain reaction was used to quantify the abundance of the CTX-M gene family – the most wide-spread ESBL gene family – and the 16S rRNA gene as a proxy for bacterial load. We find strong dynamic heterogeneity in CTX-M abundance that is largely explained by the variable nature of the swab sampling. Using information on time-varying antibiotic treatments, we develop a dynamic Bayesian model to decompose the serial data into observational variation and ecological signal and to quantify the potentially causal antibiotic effects.

We find an association of treatment with cefuroxime or ceftriaxone with increased CTX-M abundance (approximately 21% and 10% daily increase, respectively), while treatment with meropenem or piperacillin-tazobactam is associated with decreased CTX-M (approximately 8% daily decrease for both). Despite a potential risk for indirect selection, oral ciprofloxacin is also associated with decreasing CTX-M (approximately 8% decrease per day). Using our dynamic model to make forward stochastic simulations of CTX-M dynamics, we generate testable predictions about antibiotic impacts on duration of carriage. We find that a typical course of cefuroxime or ceftriaxone is expected to more than double a patient’s carriage duration of CTX-M. A typical course of piperacillin-tazobactam or of meropenem – both options to treat hospital acquired infections (HAI) like pneumonia – would reduce CTX-M carriage time relative to ceftriaxone plus amikacin (also an option to treat HAIs) by about 70%. While most antibiotics showed little association with changes in total bacterial abundance, meropenem and piperacillin-tazobactam were associated with decrease in 16S rRNA abundance (3% and 4% daily decrease, respectively).

Our study quantifies antibiotic impacts on within-host resistance abundance and resistance carriage, and informs our understanding of how changes in patterns of antibiotic use will affect the prevalence of resistance. This work also provides an analytical framework that can be used more generally to quantify the antibiotic treatment effects on within-host dynamics of determinants of antibiotic resistance using clinical data.

## Introduction

Antibiotic use can select for resistance through multiple pathways (Lipsitch & Samore, 2002): It may i) affect the duration of resistance carriage and hence transmission potential, ii) increase bacterial load of resistant organisms and thus increase transmission, or iii) selectively suppress host microbial flora where resistance is lacking, which may reduce the potential for transmission of sensitive organisms and also render those people more susceptible to cross-infections with resistant bacteria. For all of these processes, a quantitative understanding of within-host selection dynamics is important.

Here we focus on Enterobacteriaceae, a bacterial family that is commonly found in the healthy mammalian gut microbiome (Donnenberg, n.d.). Some member genus-species—*Klebsiella pneumonia*, *Escherichia coli*, *Enterobacter spp*.—are important opportunistic human pathogens that can cause urinary tract, bloodstream, and intra-abdominal infections, as well as respiratory tract infections such as hospital-acquired pneumonia. A major concern is the global increase in extended-spectrum beta-lactamase (ESBL)-producing organisms in this family (Tacconelli et al., 2018). ESBL genes – of which the most important and globally widespread is the bla_CTX-M_ gene family – confer resistance to clinically important broad-spectrum antimicrobials, such as third generation cephalosporins (D. L. Paterson, 2000). These genes commonly reside on large conjugative plasmids (Bonnet, 2003), and are co-carried with other antibiotic resistance determinants, making them a good marker for multi-drug resistance (MDR) in the Enterobacteriaceae (Schwaber, Navon-Venezia, Schwartz, & Carmeli, 2005). Because Enterobacteriaceae have their main biological niche in the gut microbiome (Masci, 2005), these bacteria are exposed to substantial collateral selection from antibiotics used to treat or prevent infections with other organisms (“bystander selection”(Tedijanto, Olesen, Grad, & Lipsitch, 2018)). Quantifying the effects of antibiotic therapy on the within-host resistance dynamics will help us to better understand the potential for selection of drug-resistant Enterobacteriaceae associated with different patterns of antibiotic usage.

In this work, we analysed follow-up rectal swabs (n=833) from 133 ESBL positive hospitalised patients from three hospitals (Italy, Romania, Serbia) to study the dynamics of antibiotic resistance gene abundance. Both CTX-M gene and 16S rRNA gene abundance, as a proxy for total bacterial load, were determined using quantitative polymerase chain reaction (qPCR). Previously, (Meletiadis et al., 2017) demonstrated a statistical association between exposure to ceftriaxone and increases in relative abundance of CTX-M using a subset of these data. Here, we aimed to fully characterise the within-host dynamics of CTX-M by developing a mechanistic model relating changes in CTX-M and 16S rRNA abundance to fine-grained patient-level antibiotic exposure data including all important classes of antibiotics used in this population. We extended previous work that uses discrete time Markov models to infer ecological parameters from microbial ecosystems (Faust & Raes, 2012; Stein et al., 2013). By incorporating hidden-state dynamics, our model explicitly accounts for observation uncertainty (due to variability in qPCR measurements and the rectal swab procedure), allowing us to separate observation noise from real within-host processes. We then used this model to make predictions about how patterns of exposure to different antibiotics would impact on CTX-M carriage duration. The development of this data-driven within-host model and its use in exploring the impact of antibiotic treatment on amplification and loss of resistance is an important step in developing a quantitative mechanistic understanding of how antibiotic use drives changes in the prevalence of resistance in a population.

## Results

### 1 Patient cohort and treatment

During enrolment time of the study a total of 1,102 patients were screened positive for ESBL Enterobacteriaceae at admission, and 133 patients (12%) gave consent to be included in the study: 51 (38%) from Romania; 52 (39%) from Serbia; and 30 (23%) from Italy. The median length of hospital stay was 15 days (maximum of 53 days), and a median of five rectal swabs per patient were taken (range of 1-15). 114 out of 133 (86%) enrolled patients received antibiotics during their stay and 85 of these, 114 (75%) received two or more different antibiotics. The different antibiotic classes, ranked by proportion of antibiotic treatment days, were cephalosporins (22.9%), antimycobacterials (18.8%), fluoroquinolones (16.3%), penicillins (8.3%), imidazole derivatives (8%), glycopeptide (7%), carbapenems (4.6%), and others (14.1%). Two thirds of antibiotic treatment days were from intravenously administered antibiotics and one third from oral administration. Details on individual antibiotics are given in **Table 2**.

**Table 1.** Summary of the study

|  |  |
| --- | --- |
| Number of participating hospitals | 3 (Serbia, Italy, Romania) |
| Study duration | 2 years (Jan 2011 - Dec 2012) |
| Inclusion criteria | inpatients of medical & surgical wards, adults, non-pregnant, ESBL *Enterobacteriaceae* carriers (at admission) |
| Number of patients followed up | 133 (including 1 with a single swab taken) |
| Intervals between rectal swabs | 2 to 3 days |
| qPCR targets | CTX-M (ESBL resistance gene), 16S rRNA (total bacterial load) |
| Number of different antibiotics used | 37 |

**Table 2.**
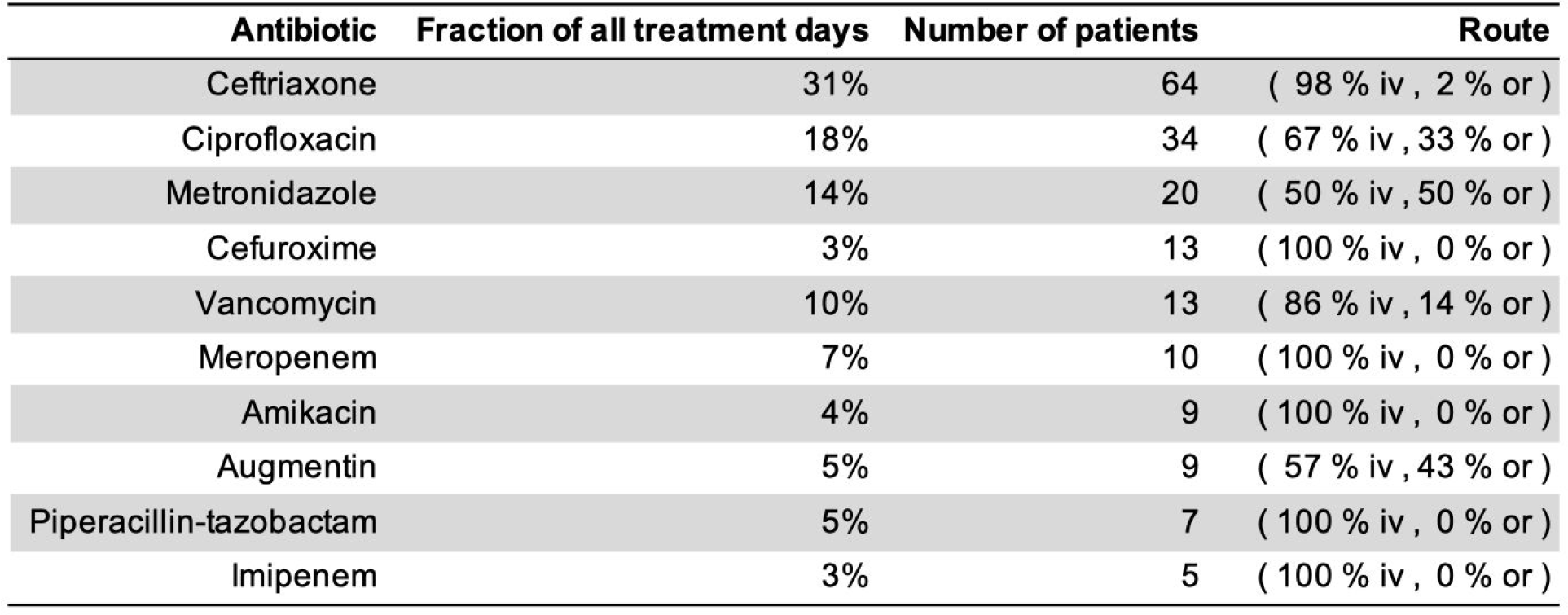
Overview of antibiotic treatments showing the ten most used antibiotics of the study. (iv: intravenous, or: oral)

| Antibiotic | Fraction of all treatment days | Number of patients | Route |
| --- | --- | --- | --- |
| Ceftriaxone | 31% | 64 | ( 98 % iv , 2 % or ) |
| Ciprofloxacin | 18% | 34 | ( 67 % iv , 33 % or ) |
| Metronidazole | 14% | 20 | ( 50 % iv , 50 % or ) |
| Cefuroxime | 3% | 13 | ( 100 % iv , 0 % or ) |
| Vancomycin | 10% | 13 | ( 86 % iv , 14 % or ) |
| Meropenem | 7% | 10 | ( 100 % iv , 0 % or ) |
| Amikacin | 4% | 9 | ( 100 % iv , 0 % or ) |
| Augmentin | 5% | 9 | ( 57 % iv , 43 % or ) |
| Piperacillin-tazobactam | 5% | 7 | ( 100 % iv , 0 % or ) |
| Imipenem | 3% | 5 | ( 100 % iv , 0 % or ) |

### 2 Resistance dynamics

The time-varying CTX-M abundance exhibits a diverse range of dynamic patterns, including monotonic increases and decreases, as well as highly variable non-monotonic behaviour (**Figure 1 a**). Qualitatively similar fluctuations in CTX-M abundance were seen both in the presence and absence of antibiotic treatment. To determine whether this high level of dynamic variation contained a meaningful biological signal, we first studied temporal autocorrelation. If the observed variability is driven by observation uncertainty – for instance through the swab procedure, DNA extraction, or qPCR process – we expect little autocorrelation in the time series. Conversely, if the observed fluctuations reflect true within-host dynamics in carriage levels, we would generally expect to see positive autocorrelation. We found a clear signal of first-order autocorrelation for both the CTX-M and the 16S rRNA gene time series, though autocorrelation was substantially stronger for the CTX-M data (**Supplementary Figure 1 a and b**). Using a Bayesian state-space model that decomposes the time series data into an observation component (representing noise due to variability in qPCR runs and in the swab and DNA extraction procedure) and a process component (due to the within-host dynamics), we estimated that much of the variability in CTX-M and 16S rRNA outcomes was due to measurement error associated with the swab procedure (median estimate [90% credible interval [CrI]] of 54% [44%, 57%] and 73% [68%, 77%], respectively). However, the CTX-M data in particular were found to contain a strong process component signal, indicating that a median estimate of 36% (90% CrI 30%, 43%) of the variability in the qPCR outcomes was due to underlying within-host dynamics (**Supplementary Figure 1 c**). To further investigate the determinants of CTX-M gene variation, we explored how much the CTX-M gene load varied between different patients or, over time, within the same patient. Using a Bayesian state-space model (see Materials and Methods) we found 16S rRNA gene abundance to be two orders of magnitude higher than CTX-M (median ratio 16S / CTX-M [90% CrI] 158 [88, 181]), with an estimated coefficient of variation (ratio of standard deviation to the mean) of 5.5 for 16S rRNA and 32.1 for CTX-M. Between-patient abundance of CTX-M showed substantially more variability than within-patient abundance (median ratio [90% CrI] 134 [18, 1422]). In contrast, 16S rRNA gene abundance had similar between-patient and within-patient variability (median ratio [90% CrI] 0.8 [0.4,1.7]) (**Figure 2**).

**Figure 1.**
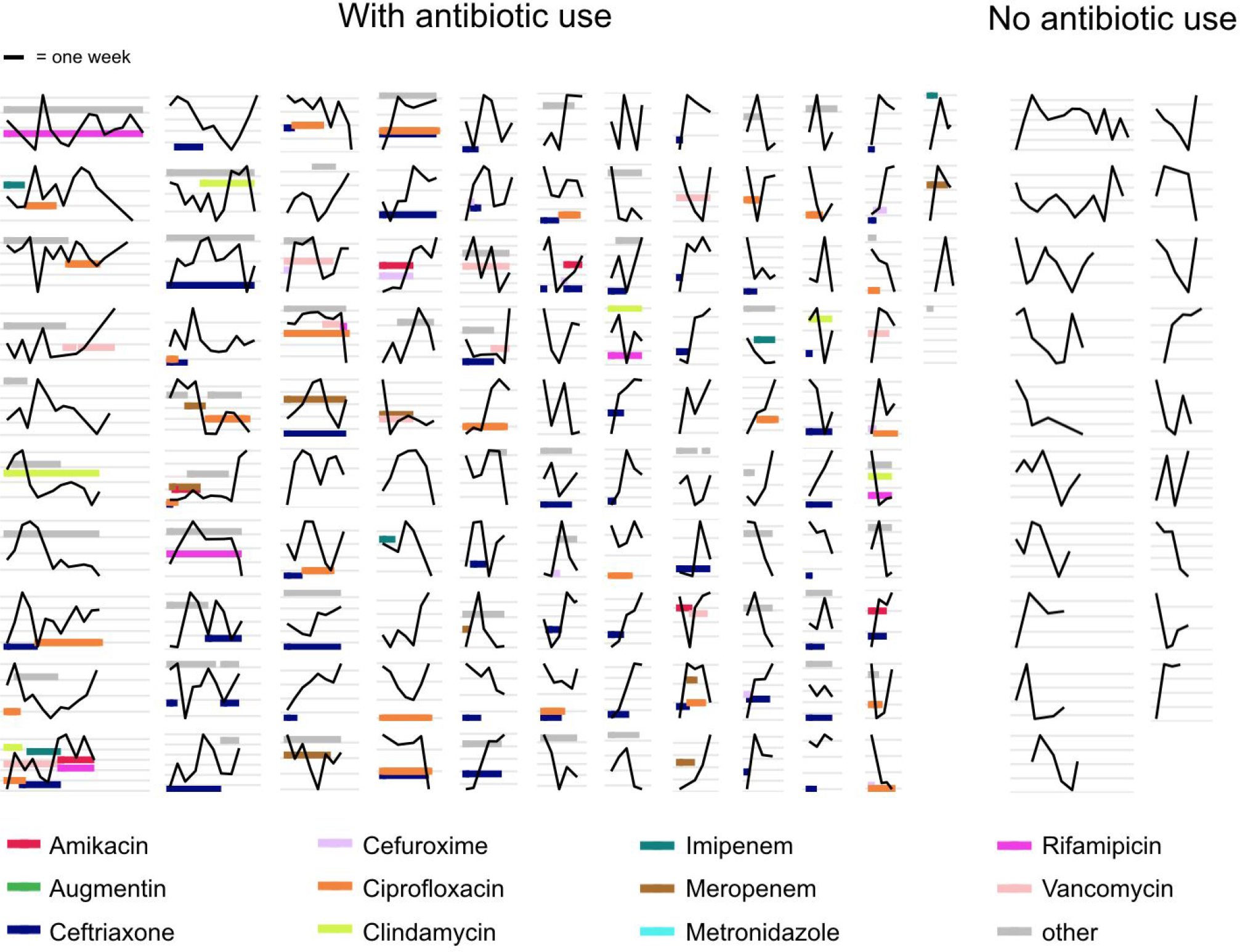
**a**) Time series plots demonstrating the diverse range of dynamical patterns of CTX-M resistance gene abundance across the 133 included patients. The x-axis scale is identical across panels, the length of one week is given for scale in the top-left corner. Timelines are ordered by length. The y-axis scale differs between panels, with the space between vertical grey lines representing a 10-fold change in the absolute CTX-M gene abundance (measured in copy numbers). The left-hand side shows patients who received antibiotic treatment (n=114), and the two right-hand side columns are patients without antibiotic treatment (n=19). For clarity, we show only the twelve most frequently used antibiotics in distinct colours and other antibiotics in grey.

**Figure 2.**
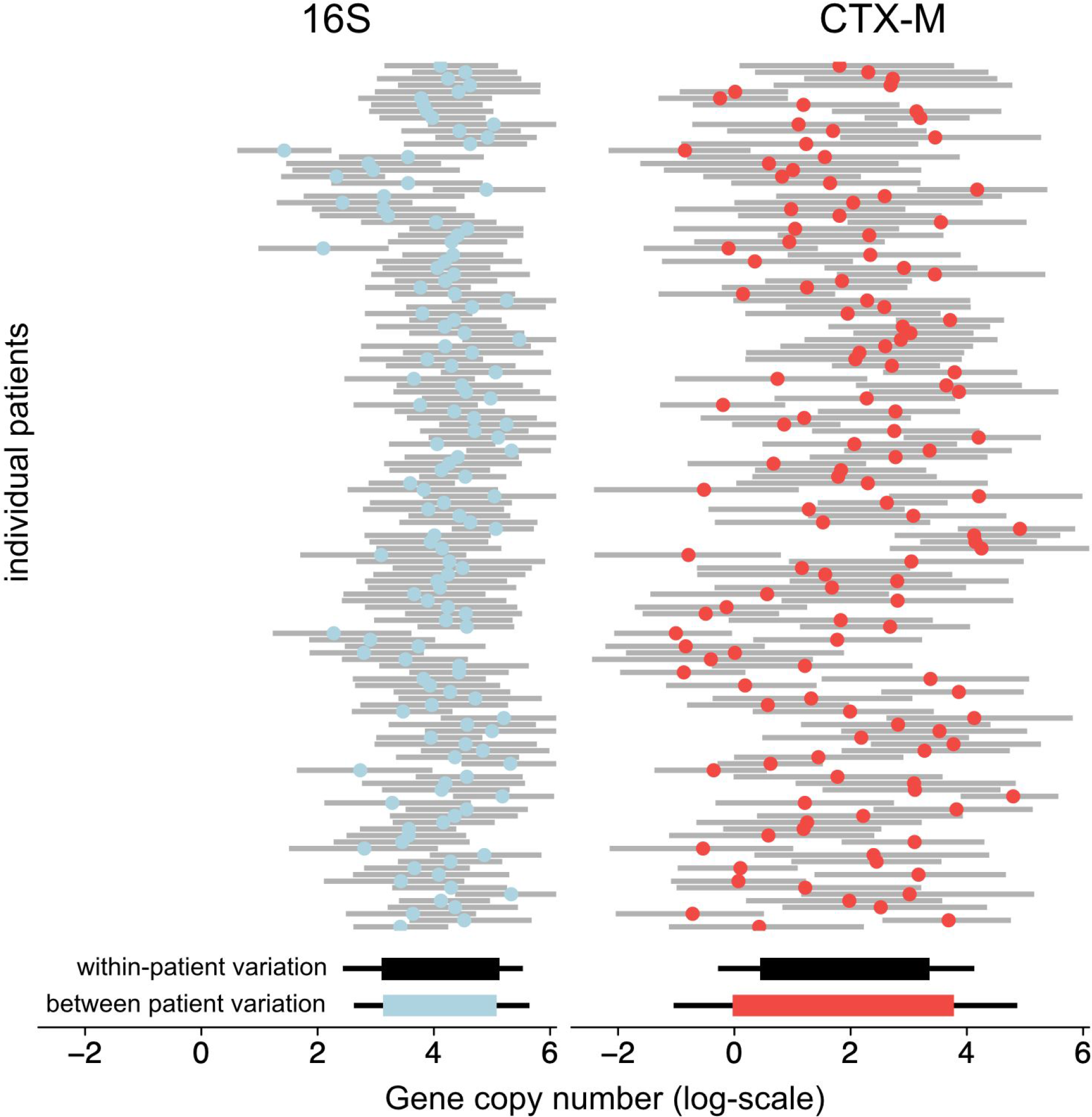
Variability of 16S abundance and CTX-M abundance for individual patients and comparison of within and between patient variation using a Bayesian hierarchical model. The upper part shows individual patient intercepts given as mean posterior estimates (colored dots) together with posterior predictions for each patient (grey bars show 80% central quantiles). The lower part shows the within-patient variation as simulated outcomes using the mean population intercept and variance (black bars), and the between-patient variation as the distribution of patient intercepts in coloured bars (thick bars and thin bars show the 80% and 95% central quantiles, respectively).

### 3 Associating resistance and antibiotic treatment

The change in relative resistance between samples, measured as CTX-M abundance divided by 16S rRNA gene abundance, was only slightly elevated in time intervals where antibiotics were given compared to those where they were not (**Figure 3 a**). However, use of antibiotics with expected activity against ESBL producers (doxycycline, ertapenem, meropenem, tigecycline, colistin, augmentin, ampicillin-sulbactam, piperacillin-tazobactam, amikacin, gentamicin, ciprofloxacin, imipenem, levofloxacin) was associated with a modest decrease in CTX-M abundance (**Figure 3 b**). In contrast, the use of antibiotics with broad spectrum killing activity but no activity against ESBL producers (amoxicillin, ampicillin, cefepime, cefotaxime, ceftazidime, ceftriaxone, cefuroxime, ofloxacin, sulfamethoxazole-trimethoprim) was associated with substantially higher increases in relative CTX-M abundance (**Figure 3 c**).

**Figure 3.**
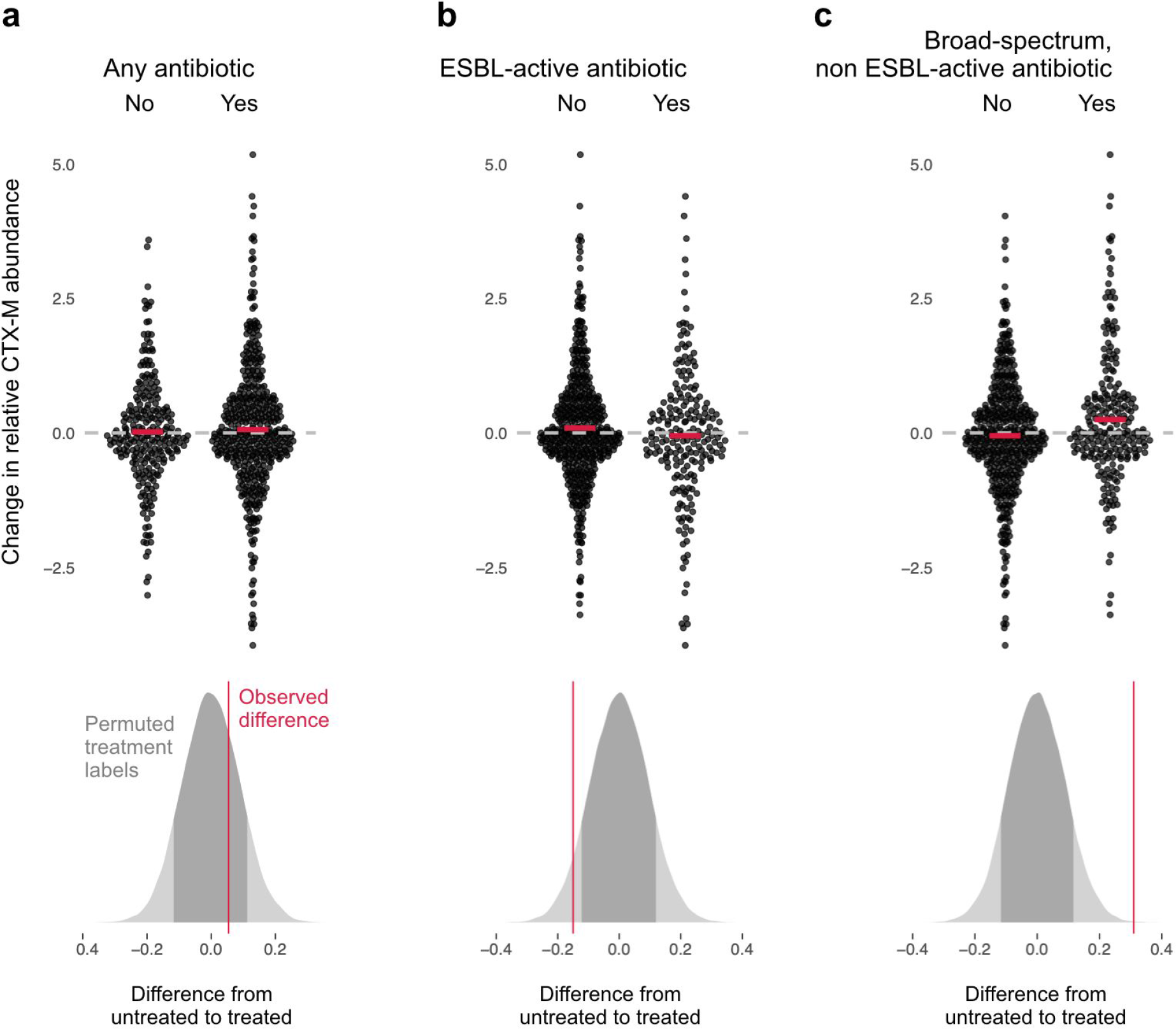
Association of antibiotic use with change in relative resistance (abundance of CTX-M divided by abundance of 16S rRNA). The upper panels show the change in relative resistance between all neighbouring timepoints (black dots), dashed horizontal lines in grey indicate the region of no change. Pairs of violin scatter plots (with the mean values shown as red bars) contrast different treatment that occurred between those timepoints, ‘Yes’ indicates treatment with specified antibiotics and ‘No’ means other or no treatment. The lower three panels show the distribution of mean differences of the change in relative resistance between treatment groups generated through treatment-label permutation (areas in darker grey show 80% central quantiles). The distributions are overlaid with the observed difference (red vertical line). Subplot **a**) compares treatment with any antibiotic versus no antibiotic. Subplot **b**) compares treatment with antibiotics with expected activity against most ESBL-producers (doxycycline, ertapenem, meropenem, tigecycline, colistin, augmentin, ampicillin-sulbactam, piperacillin-tazobactam, amikacin, gentamicin, ciprofloxacin, imipenem) with all other treatment, including no treatment. Finally, in subplot **c**) we consider antibiotics with broad-spectrum activity but mostly lack of activity against ESBL producing strains (amikacin, amoxicillin, ampicillin, cefepime, cefotaxime, ceftazidime, ceftriaxone, cefuroxime, levofloxacin, ofloxacin, sulfamethoxazole-trimethoprim).

### 4 Mechanistic antibiotic effect model

Fitting a dynamic model of CTX-M abundance and 16S rRNA abundance to the data, we found that cefuroxime and ceftriaxone were associated with increases in both absolute CTX-M abundance (mean daily increase [90% CrI] 21% [1%, 42%] and 10% [4%, 17%], respectively) and relative CTX-M abundance (14% [-1%, 30%] and 11% [5%, 17%], respectively) (**Figure 4**). Piperacillin-tazobactam, meropenem and ciprofloxacin (when given orally) were negatively associated with both CTX-M (−8% [-18%, 2%], −8% [-17%, 1%], and −8% [-17%, 2%], respectively) and 16S rRNA gene abundance (−3% [-8%, 1%], −2% [-7%, 1%], and −1% [-6%, 3%], respectively), although uncertainty is large (**Figure 4**). Their effect on relative resistance (CTX-M / 16S rRNA) also appears to be negative (−5% [-14%, 5%] for piperacillin-tazobactam, −5% [-14%, 4%] for meropenem, −7% [-15%, 3%] for oral ciprofloxacin). Intravenously administered ciprofloxacin did not show these effects. Imipenem and meropenem had similar effects on CTX-M abundance, while no clear effects were evident for amikacin, metronidazole, and augmentin. With the dynamic model, we are able to make predictions about the time required for the CTX-M genes to fall below detection levels. To achieve this, we add to our stochastic model a threshold below which the CTX-M gene cannot be detected (see *Materials and Methods*). The predictions show a high degree of uncertainty, visible as long-tailed predictive distributions. Because of the skew, we report here the median instead of the mean together with 80% credible intervals. We find that a single 8-day course of cefuroxime or a 14-day course of ceftriaxone substantially prolongs carriage of CTX-M, by a median estimate of 147% (80% CrI 13.4%, 577%) for cefuroxime and 120% (80% CrI −8.6%, 492%) for ceftriaxone versus no exposure (**Figure 5, upper panel**). Addition of oral ciprofloxacin to a course of augmentin or ceftriaxone reduces CTX-M carriage duration (by approximately 51% [80% CrI −115%, 89%] and 48% [80% CrI −71.1%, 86%]) (**Figure 5, upper panel**). A typical 14-day course of meropenem or a 8-day course of piperacillin-tazobactam reduce CTX-M carriage time relative to no treatment (by approximately 42% [80% CrI −25%, 75%] and 41% [80% CrI −45%, 71%], respectively), and each reduces CTX-M carriage even more relative to a 7-day course of combined ceftriaxone plus amikacin (by approximately 69% [80% CrI 20%, 89%] and 66% [80% CrI −7%, 88%], respectively) (**Figure 6, middle panel**). Finally, a 14-day course of meropenem reduces CTX-M resistance carriage relative to a shorter 5-day course (by approximately 69% [80% CrI 20%, 89%]) (**Figure 6, lower panel**).

**Figure 4.**
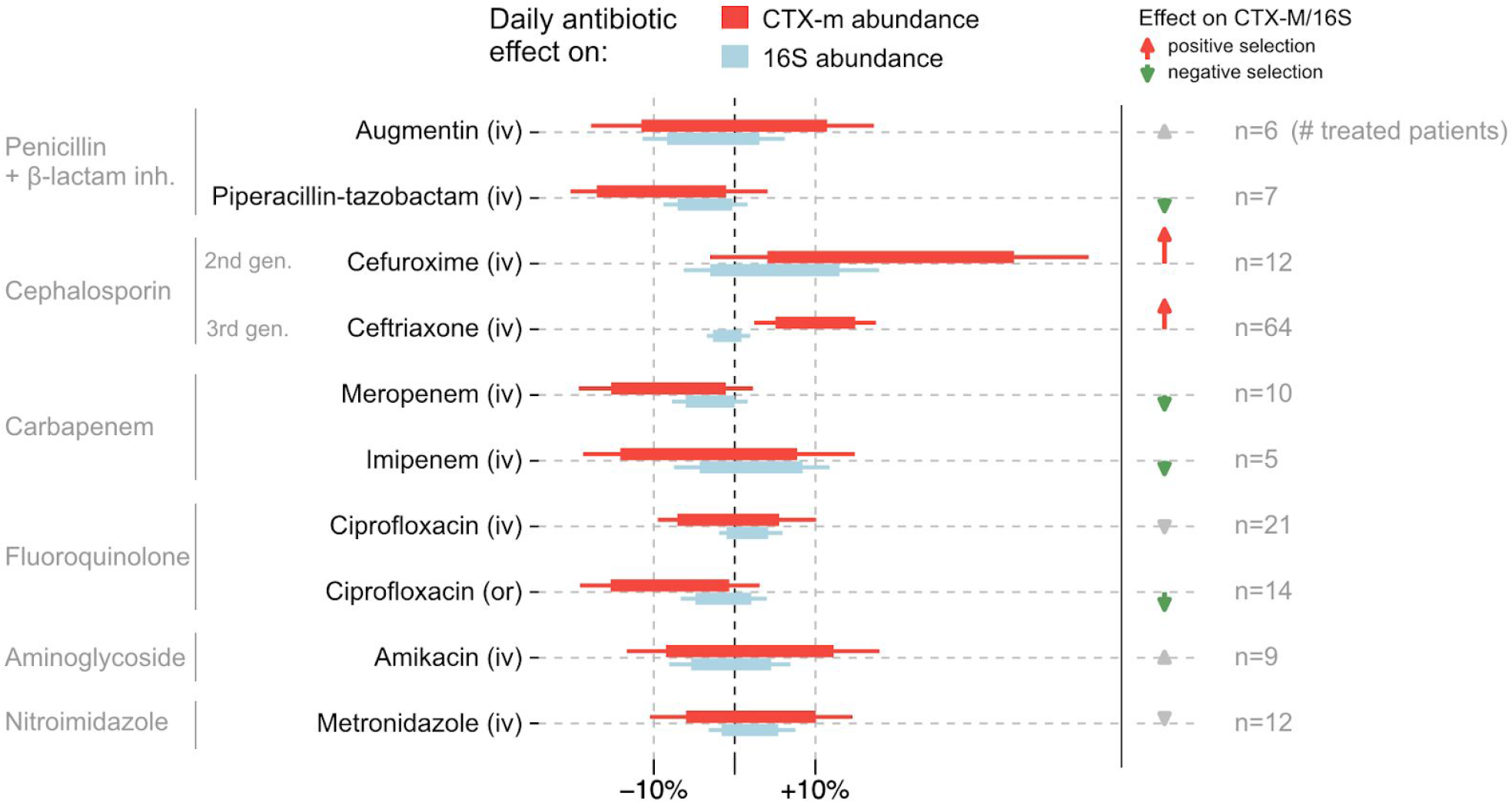
Estimated effects of different antibiotics on within-host dynamics from a multivariable model. The bars show estimated daily effects of individual antibiotics on the absolute CTX-M abundance (red) and 16S rRNA abundance (light blue) indicating the 80% and 95% highest posterior density intervals (thick and thin horizontal bars, respectively). The model also gives the antibiotic effect on the CTX-M / 16S relative resistance shown as arrows on the right-hand side. Arrows are in grey for antibiotics with mean effect estimates between −10% and +10%, otherwise they are coloured red (positive selection) and green (negative selection).

**Figure 5.**
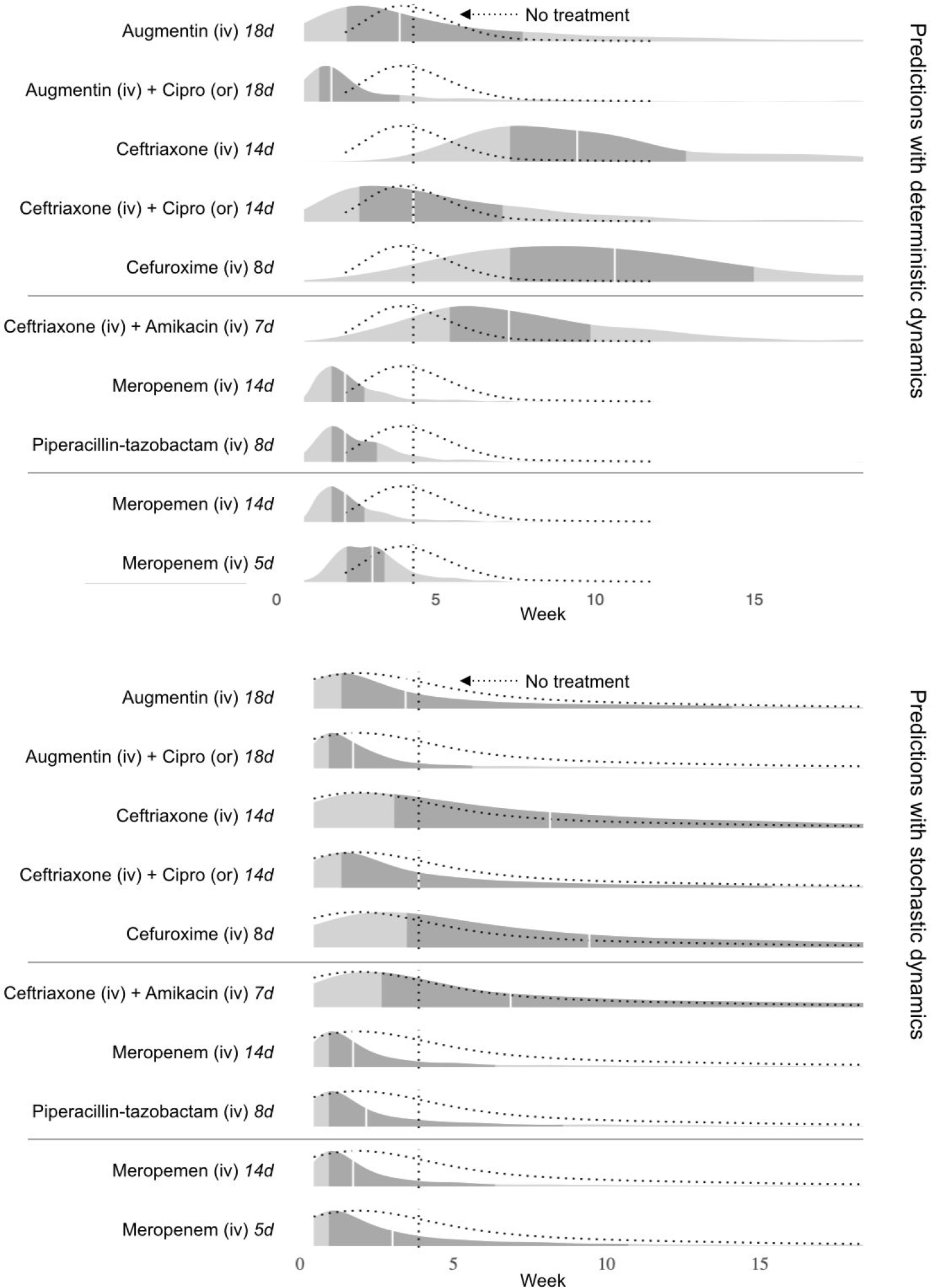
Simulated predictions of CTX-M carriage duration under different alternative antibiotic treatments. The upper panel shows model predictions with parameter uncertainty, but assuming deterministic dynamics. The lower panel shows the predictions with parameter uncertainty as well as Markov process uncertainty. The darker grey areas shows the 50% credible intervals and the white lines show the median predictions. Both panels compare the same treatments. Each density distribution is overlaid with the density line of the no treatment case (dotted line) and its median prediction (dotted vertical line). We compare predictions for treatment with augmentin (18 days) and ceftriaxone (14 days), and each in combination treatment with ciprofloxacin. We also compare treatment with ceftriaxone plus amikacin (7 days), meropenem (14 days), and piperacillin-tazobactam (8 days). Finally, we compare a normal course of meropenem (14 days) with a shortened course (5 days).

## Discussion and Conclusion

By fitting a dynamic model accounting for both observation noise and within-host dynamics to time series data from 133 patients, we quantified the association between antibiotic exposure and changes in rectal swab abundance of gut bacteria and CTX-M resistance genes. The largest effects were found for exposures to the second and third generation cephalosporins cefuroxime and ceftriaxone, both of which were associated with increases in CTX-M abundance. Forward simulations indicated that if these associations are causal, exposure to typical courses of these antibiotics would be expected to more than double the expected carriage duration of CTX-M. Both cefuroxime and ceftriaxone have broad-spectrum killing activity (McLeod, Nahata, & Barson, 1985; Neu & Fu, 1978), but have limited activity against ESBL-producing organisms (Livermore & Brown, 2001; Sorlózano et al., 2007). Therefore, a direct selective effect is biologically plausible to account for the above finding. Surprisingly, despite the relatively broad antibacterial spectrum of cefuroxime and ceftriaxone, there was no evidence that exposure to these antibiotics reduced 16S rRNA abundance. This may reflect high parameter uncertainty, or that killed bacterial strains may quickly be replaced by overgrowth of other, non-susceptible, strains (Hildebrand et al., 2019). Further, 16S abundance as measured by qPCR may not be a good proxy for the abundance of living bacterial cells. When bacteria are killed by antibiotics, we expect that shedding of dead cells from the gut could delay a measurable reduction in 16S rRNA or even lead to a temporary increase. It is therefore possible that we underestimated the effects of all antibiotics on suppressing the bacterial load.

Though credible intervals were wide, meropenem, piperacillin-tazobactam, and oral ciprofloxacin were all associated with reductions in CTX-M abundance. All three are broad-spectrum antibiotics with activity against ESBL producers in the absence of specific co-resistance, and are common agents for treating hospital-acquired infections (Lautenbach et al., 2001; Masterton, Drusano, Paterson, & Park, 2003; David L. Paterson, 2006). They were also associated with a negative effect on relative resistance, CTX-M divided by 16S rRNA gene abundance. This observation can be explained by a general reduction of bacterial biomass that leads the CTX-M abundance to drop below detection levels. In line with this, simulations suggest that a typical course of meropenem or of piperacillin-tazobactam would reduce CTX-M carriage time relative to no treatment by about 40%, and each reduces CTX-M carriage duration by about 70% relative to a combined course of ceftriaxone plus amikacin. Also, a standard (14 day) course of meropenem is found to reduce ESBL resistance carriage relative to a shortened course (5 day) by approximately 70%. While these findings suggest suppression of ESBL producing bacteria by meropenem, it may at the same time increase selection for carbapenem-resistance. Finally, we also find that adding oral ciprofloxacin to augmentin or ceftriaxone reduces ESBL-producing bacteria carriage by approximately 50%. This is slightly surprising, because CTX-M carrying bacteria commonly co-carry resistance to other antibiotics in bacteria, especially resistance to ciprofloxacin, which would lead to indirect selection of ESBL through other antibiotics. In this study population, there appears to be little resistance to meropenem or to ciprofloxacin, as indicated by the CTX-M suppressing effect of these antibiotics. The effects of meropenem and ciprofloxacin on CTX-M resistance could vary depending on the local epidemiology of carbapenem and fluoroquinolone resistance. Although oral ciprofloxacin showed an association with reduced CTX-M abundance, intravenous ciprofloxacin showed near zero effect. Antibiotic selection for resistance with different routes of administration has been previously explored in a mouse model, which suggested that oral drug administration has stronger selective effect on resistance than intravenous administration (Zhang et al. 2013), but similar studies for humans are lacking. Delineating the relationship between the various routes of antibiotic administration and resistance, as we do here, will be important for informing stewardship guidelines about switching between intravenous and oral therapy.

Few previous studies have quantified the association between antibiotic use and resistance abundance in the human microbiome. Two studies involving patients admitted to intensive care units looked at the effect of a preventative antibiotic cocktail (selective digestive decontamination) on microbiome resistance in patients, with one study (including n=13 patients) finding no clear effect (Buelow et al., 2014), and the other (n=10) showing increases of four different resistance genes associated with treatment (Buelow et al., 2017). These studies did not attempt to model patient time-series since only a maximum of three time-points per patient were sampled. Compared to these studies, we considered here a 10-fold bigger patient cohort of 133 patients and with a total of 833 time points, a median of 5 per patient. We are not aware of any other data-driven mechanistic model of the relationship between antibiotic exposure and resistance gene abundance in the human gut, and we believe that this presents a useful analytical framework that can be adapted in the context of other studies to quantify the antibiotic treatment effects on within-dynamics from clinical data. Abundance of resistance measured in a patient’s stool has been shown to be predictive of the risk of the patient getting a resistant infection (Ruppé E, n.d.; Woerther et al., 2015). Thus predictive models, such as ours, can help to identify antibiotic exposures that would minimise this risk. Another major benefit of the modelling framework we have developed is the ability to make testable predictions about the impact of different antibiotics on the duration of carriage of resistant determinants at levels above detection thresholds. Understanding this impact of antibiotics on carriage duration is a key step in developing a mechanistic understanding of the relationship between the frequency with which antibiotics are used in a population and the proportion of the population in whom resistance can be detected. Such an understanding relies on quantifying the antibiotic effects in individual exposed patients, as we do here, but also on quantifying the knock-on effects on transmission to contacts. These indirect effects are likely to be considerable. A recent study in Dutch travellers returning to the Netherlands who had acquired ESBL carriage overseas found that their new ESBL carriage status was associated with a 150% increase in the daily risk of non-carrying household members becoming ESBL positive (Arcilla et al., 2017). Developing mechanistic models for the spread of ESBLs and other resistance determinants within host populations accounting for such direct and indirect antibiotic effects is an important priority for future research. Such models would help us to understand and predict how changes in antibiotic usage patterns affect the prevalence of antimicrobial resistance in a community and would help to prioritise interventions to reduce the burden of antimicrobial resistance.

## Materials and Methods

### Study participants and follow-up

This was an observational, prospective, cohort study that included data from three hospitals (Italy, Serbia and Romania), with known high prevalence rates of antibiotic resistance in bacterial infections. The study was conducted over two years from January 2011 to December 2012 as part of the multi-centre SATURN (‘Impact of Specific Antibiotic Therapies on the prevalence of hUman host ResistaNt bacteria’) project (NO241796; clinical trials.gov NTC01208519). The study enrolled adult (>18 y) inpatients of medical and surgical wards, excluding pregnant patients. Enrolled patients were screened at admission for carriage of ESBL-producing *Enterobacteriaceae* with rectal swabs (E swab, Copan, Italy). Patients who tested positive for ESBL carriage (details below) were included in the follow-up cohort. For all follow-up patients (n=133) rectal swabs were taken every two to three days during hospitalisation, which includes one swab at admission and one at discharge. The swabs were stored at −80 degrees Celsius and sent to a central laboratory for processing. The study also collected information on antibiotics administered, duration, and route of administration. See **Table 1** for an overview of the study details.

### ESBL identification

Samples taken at admission were cultured on chromogenic agars (Brilliance ESBL, Oxoid, Basingstoke, UK) to test for ESBL positive organisms. ESBL status was confirmed with the double disk diffusion method.

### Quantitative PCR

DNA was extracted from the swab samples and a fixed volume of DNA solution was used as a template for quantitative PCR (qPCR) assays. Two singleplex qPCR assays were conducted, one to assess quantity of CTX-M gene family with primers CTX-M-A6 (TGGTRAYRTGGMTBAARGGCA) and CTX-M-A8 (TGGGTRAARTARGTSACCAGAA) (product length, 175 bp) and one targeting a conserved bacterial 16S rRNA gene region bacteria using the following primer set, 16S_E939F (GAATTGACGGGGGCCCGCACAAG) and 16S_1492R (TACGGYTACCTTGTTACGACTT) (product length, 597 bp) to assess total bacterial quantity. Quantification with qPCR was carried out mostly in duplicates, with some triplicates.

### Time series autocorrelation

We first transformed all qPCR measurements onto log-scale. For all patients and each time point we then computed the mean of the qPCR duplicates (or triplicates) for CTX-M and 16S rRNA. To get reliable estimates of autocorrelation, we selected only patients with more than five time points. Separately for the CTX-M and 16S rRNA gene data, we computed the first-order autocorrelation (disregarding varying spacing between time points) for each patient, and we averaged these values across the patients. We then simulated serially uncorrelated “white noise” time series, again separately for CTX-M and 16S rRNA, with the same length as the patient data and with identical time series mean and variance. Similar to the real data, we computed mean autocorrelations for the simulated data and show their distribution for a large number of simulations (n=10,000) together with the observed autocorrelation (**Supplementary Figure 1 a and b**). We also computed the proportion of simulated datasets that showed an average autocorrelation equal to, or larger than, the observed data, and we show those numbers on the arrows in **Supplementary Figure 1 a and b**.

### Estimating observation and process noise in time series

To estimate the amount of observation noise and process noise in the time series we constructed a Bayesian state-space model that included qPCR noise, swab noise, and biological noise. This model is given through:

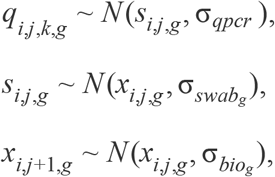

where *i* denotes a given patient, *j* denotes a swab (one per time point), *k* denotes a qPCR measurement (multiple repeats per swab), and *g* denotes the genetic target, either CTX-M or 16S rRNA. The term *q*_*i,j,k,g*_, represents the measured quantity of genetic targe *g*, of the *k*th qPCR replicate (on a log-scale) from patient *i*, at time point *j*. In addition, there are two hidden-state parameter vectors: *s*_*i,j,g*_ is the underlying, true sequence abundance genetic target *g* that a qPCR assay with 100% efficiency could (in theory) measure at time point *j* for patient *i*, and *x*_*i,j,g*_ is the actual gene abundance of genetic target *g*, in the swab at time point *j* for patient *i*, before the added noise through the swab process and gene extraction. The unobserved variables of interest are σ_*qpcr*_, the qPCR machine error (assumed to be the same for CTX-M and 16S rRNA), 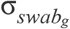, the swab variation of the genetic targe *g*, and 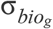, the variation of genetic target *g* from biological processes. On all parameters we assigned uniform (“flat”) priors over their legal range of values, which was [−∞, +∞] for hidden-state parameters, and [0, +∞] for the noise parameters. We then fitted this model to the CTX-M and 16S rRNA measurements. The posteriors are shown in **Supplement Figure 1 c**, where we expressed each type of noise as a fraction of the total noise. The model was fitted using Stan software (Carpenter et al., 2017) and with additional analysis in R (R Core Team, 2016), and we sampled from the posterior with four chains with 2000 iterations that included a burn-in period of 300 iterations.

### Between and within time series variance

For the estimation of between and within time series variation we used a Bayesian hierarchical model, which accounted for unbalanced sampling between patients. This model used the mean estimates of *x*_*i,j*_ (actual gene abundance in time point *j* for patient *i*) from the previous model, and it took the for

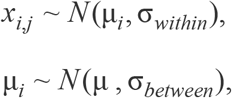

where μ_*i*_ is the mean abundance of patient *i*, around which the measurements were assumed to be normally distributed with standard deviation σ_*within*_, the within time series variation. The mean abundances were assumed to follow a normal distribution with a population mean μ and σ_*between*_, the between patient variation. We assigned uniform priors over the range of [−∞, +∞] for the population and the patient means, and over the range of [0, +∞] for the standard deviations. We fitted the model using Stan, with 4000 iterations and a burn-in period of 500 iterations. Model estimates are shown in **Figure 3**. To calculate the coefficient of variation for the non log-scaled CTX-M and 16S rRNA measurements, we use the transform described by Koopmans et al.: 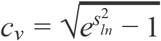 where *s*_*ln*_ is the estimated standard deviation of the log-scaled data.

### Association of antibiotic treatment and changes in resistance

To study the association between antibiotic treatment and resistance we looked at relative abundance of resistance (CTX-M abundance / 16S rRNA gene abundance) as a marker of natural selection. First, we computed the changes in relative resistance for every pair of adjacent time points and for each antibiotic we used a binary variable indicating whether or not a given antibiotic was administered between these time points. When an antibiotic treatment was on the same day as a swab, this treatment was allocated to the time interval between this day and the next swab. We first looked at how changes in relative resistance are associated with courses of any antibiotics, then with courses of antibiotics that are active against ESBL-producing bacteria (doxycycline, ertapenem, meropenem, tigecycline, colistin, augmentin, ampicillin-sulbactam, piperacillin-tazobactam, amikacin, gentamicin, ciprofloxacin, imipenem), and finally with antibiotics that have a broad-spectrum activity but no activity against ESBL (amikacin, amoxicillin, ampicillin, cefepime, cefotaxime, ceftazidime, ceftriaxone, cefuroxime, levofloxacin, ofloxacin, sulfamethoxazole-trimethoprim). Results are shown in **Figure 4**, **upper panel**. We evaluated how likely the observed differences between treatments are under the assumption of no association between treatment and resistance. For this we did a permutation or “reshuffling” experiment: we randomly reassigned (without replacement) the antibiotic treatment labels to the data intervals. We compute the distribution of mean differences from 50 000 permutations and compare this to the observed difference (**Figure 4, lower panel**).

### Mechanistic within-host model

We extended previous approaches of extracting ecological parameters from microbial ecosystem dynamics (Faust & Raes, 2012; Stein et al., 2013) by applying a Bayesian state-space model, that allowed separating process noise from observation noise and accounting for different spacing between time steps. Under the assumption that 16S rRNA gene abundance is independent of antibiotic treatment, variation in 16S rRNA would be caused mainly by the swab procedure, and it could be used to normalise CTX-M abundance. However, as we found in **Figure 5**, 16S rRNA abundance was associated with certain antibiotic treatment. Thus, we used a dynamic model that explicitly modelled antibiotic effects on 16S rRNA and on CTX-M/16S, from which the effects on CTX-M could then be computed. Studying the standard deviation between qPCR measurement repeats as a function of the mean, we observed that qPCR variation remained relatively stable over five orders of magnitude of the mean measurement (from 1.5 to 6.5 on the log scale), but it increased quickly for lower magnitudes (**Supplementary Figure 2**). In the Bayesian model for different sources of variation described above, the parameter σ_*qpcr*_ assumed that the qPCR uncertainty is the same across measurements. Here, we aim to account for the fact that low measurements of gene copy numbers have higher uncertainty. We fitted a smooth spline (choosing five degrees of freedom) to the qPCR measurements (red line in **Supplementary Figure 2**). This let us assign an estimated qPCR standard deviation to every set of qPCR repeats. We provided those estimates as data to the Bayesian model.

Our model then took the form:

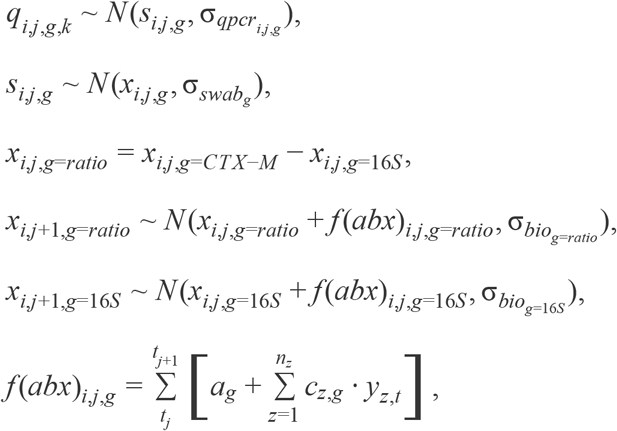

where the data is given through *q*_*i,j,g,k*_, the *k*th qPCR result (log-scaled) of patient *i*, at time point *j*, and genetic target *g* (CTX-M, 16S rRNA, or CTX-M/16S ratio), through 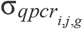, the estimated qPCR standard deviation of that set of measurements (of genetic target *g*, patient *i*, and time point *j*), and through *y*_*z,t*_, a boolean variable indicating whether or not antibiotic *z* was given on day *t*. The hidden-state variables are *s*_*i,j,g*_, the underlying, true sequence abundance at time point *j* for patient *i*, and genetic target *g*, and *x*_*i,j,g*_, the actual abundance of *g* in the swab at time point *j*, for patient *i*. The swab variability of genetic target *g* (CTX-M or 16S rRNA) is given through σ_*swab*,*g*_, and the biological variability of the CTX-M/16S ratio and of 16S are given through 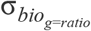 and 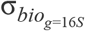. Taking the difference between the abundance of CTX-M and 16S rRNA gene yields *x*_*i,j,g*=*ratio*_, the log-scaled relative resistance of patient *i* at time point *j*. We then model the ecological dynamics with *f* (*abx*)_*i,j,g*_, the antibiotic-mediated change of *g* (either the relative resistance or 16S rRNA gene), at time point *j*, of patient *i*. We assume that changes in the relative resistance and in 16S are driven by a neutral trend (describing the dynamics in the absence of antibiotics) and by antibiotic effects. We further assume that effects are multiplicative. For example, consider a genetic target on day 0 of *x*_*t*=0_ with a neutral trend of *a*_0_. Suppose two antibiotics are given with effects *c*_1_ and *c*_2_, then on day 1 the genetic target becomes *x*_*t*=1_ = *x*_*t*=0_ · *a*_0_ · *c*_1_ · *c*_2_. Thus, the ecological dynamics term loops over the days from *t*_*j*_, the day of swab *j*, until *t*_*j*+1_, the day of the following swab of the same patient *i*, and it sums the neutral effect *a*_*g*_ and the antibiotic effect, which is given as product of effect of antibiotic *z* (*c*_z,*g*_) and the boolean indicator for treatment with *z* on day *t* ( *y*_*z,t*_), summed over all antibiotics *n*_*z*_. Note, that summing the effects of *a* and *c* for dynamics on a log-scale is equivalent to multiplicative effects on the original scale. On the hidden-state variables we assigned uniform priors over the range [−∞, +∞], on the standard deviations describing swab and biological variability we assigned uniform prior over the range [0, +∞], and on the antibiotic effects (*c*) we assigned conservative priors of the form N(0, 0.1). We fitted the model using RStudio and Stan software, and we sampled form the posterior with four parallel chains of 9000 iterations, which included a warm up phase of 2000 iterations.

We forward simulated CTX-M data using the dynamical model above and the posterior distributions from the model fit. We added to the model a threshold below which the CTX-M gene becomes extinct or at least undetectable. According to a study of returning European travelers to Southeast Asia, ESBL carriers lose their resistant bacteria after a median of 30 days (Arcilla et al., 2017). Accordingly, we simulated CTX-M time series without antibiotic treatment and chose an extinction threshold (0.25 CTX-M copy numbers) that achieved the same median extinction time. We then used this model to repeatedly (2,000 times) simulate CTX-M carriage times, with each simulation using a new draw from the parameter posterior. The resulting distribution of carriage times contains both the uncertainty in the parameter estimates and uncertainty from the Markov process. We also simulated CTX-M carriage times repeatedly (200 times) with a single set of parameter values from the posterior, then taking the median carriage time which removes Markov process uncertainty, before drawing a new set of parameters and repeated this procedure 300 times. We used both of the above methods to simulate carriage time under different alternative antibiotic treatment. The resulting distributions are shown in **Figure 6**.

### Code and data availability

R code and Stan code for the data analysis described above as well as the data will be made available online.

## Acknowledgements

We thank Jonas Schluter, Marc Lipsitch, and Thomas Crellen for valuable feedback along the way. RN as well as the study were supported by funding from the European Community’s R-GNOSIS Integrated project (FP7/2007-2013) under grant agreement number 241796. RN and BSC were also supported by The Medical Research Council and Department for International Development (grant number MR/K006924/1). BSC works within the Wellcome Trust Major Overseas Programme in SE Asia (grant number 106698/Z/14/Z).

---

**Supplementary Figure 1.**
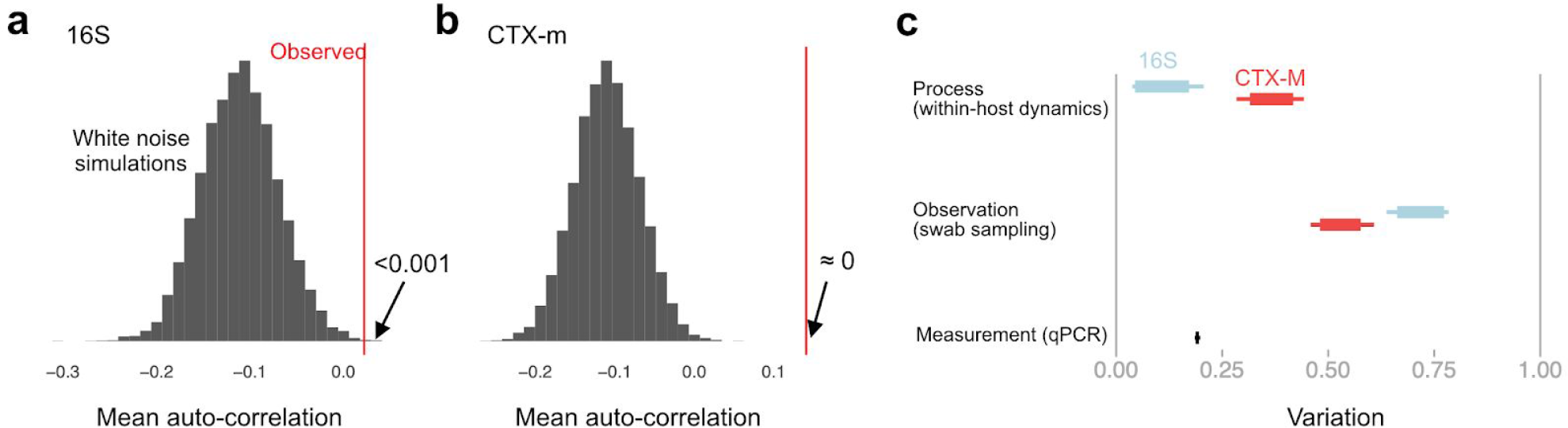
Autocorrelation and sources of variability in qPCR time series data. First-order autocorrelation averaged across all patients with more than five time points for 16S (**a**) and CTX-M (**b**) abundance data (red vertical line). This is compared to a histogram of the mean autocorrelation from 10,000 replicates of simulated serially uncorrelated “white noise” time series, simulated using the same number of observations per patient as in the real data and the same per patient mean and variance as in the observed CTX-M and 16S rRNA data. The shift toward negative autocorrelation in the simulations is an artefact due to short time series, what is of interest is the position of the observed value relative to the simulated distribution. Both CTX-M and 16S data show a clear autocorrelation signal, though autocorrelation is substantially stronger for the CTX-M data. Panel (**c**) shows Bayesian estimates of the sources of variability in the measurements, given as proportion of total variability.

**Supplementary Figure 2.**
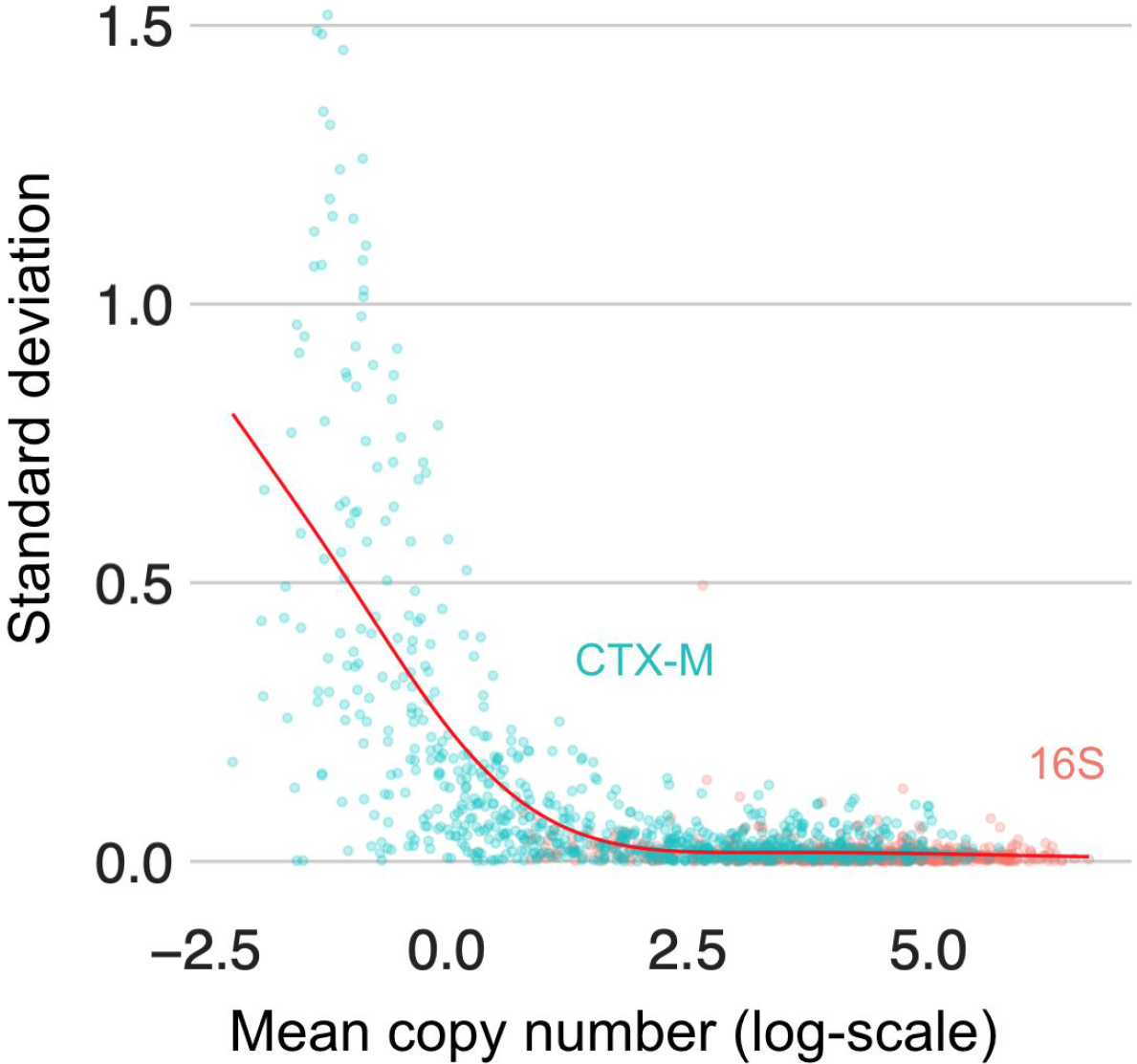
Variability in replicate qPCR runs. The standard deviation of repeat qPCR machine runs versus their mean for 16S (red) and CTX-M (turquoise). The red line represents a smooth spline fit to the data with five degrees of freedom.

